# Investigating Biomechanical Properties of *A. tortilis* as Related to their Habitat: A novel approach to characterize bending

**DOI:** 10.1101/601534

**Authors:** Ahmed Faris, Rani Altoum, Iyad Salameh, Mohammed Salameh, Saif Al-Mohannadi, Abdulla Mahmoud

## Abstract

The biomechanical properties of *Acacia tortilis* were investigated considering its habitat (wild vs. nursery). The plant materials were collected from partially urbanized area in Doha city and from Qatar Foundation Nursery. The results show that *Acacia* grown in field are more flexible than those grown in nursery. Young’s Modulus of Elasticity was found to be 191 MPa and 617 MPa and the Flexural Modulus was found to be 49 MPa and 575 MPa and the breaking force was found to be 210 and 550 N for nursery and field Acacia, respectively. The deflection angle was measured using sensitive flex sensors connected to Arduino boards and was found to be higher for field Acacia (50∘). Image processing techniques were used to mathematically describe the branch motion versus time diagrams.

The plant part being investigated was covered with red tape and videotaped while subjected to a force causing it to bend. The stem was divided into 745 successive points and the change in their position with time taken frame by frame was converted into a change in position expressed through mathematical parameters. The bending movement of the branch was found to follow a power function *H* = (4001 − *e*^0.06*m*^).

**Highlight:** Field grown *Acacia* have higher values of Young’s and Flexural Moduli than nursery grown ones thus conferring them more elasticity and flexibility.

## 1 INTRODUCTION

*Acacia* spp. are native shrubs in the Middle East and African Savanna. They occupy a wide range of habitats including the tropical and subtropical areas. *Acacia tortilis* (fig. 1), is a kind of spiny shrubs that belongs to the Fabaceae family (peas). It can grow very well in extremely arid conditions, in sunny, high temperature and dry environments as well as in drained alkaline sandy soil. *Acacia* has a high economic importance due to its wide range of uses. It is used in gardening as ornamental plant and as wind breaks. *Acacia* wood is used in furniture industries. Moreover, different chemical compounds isolated from *acacia* have medical and pharmaceutical applications.

**Figure 1.**
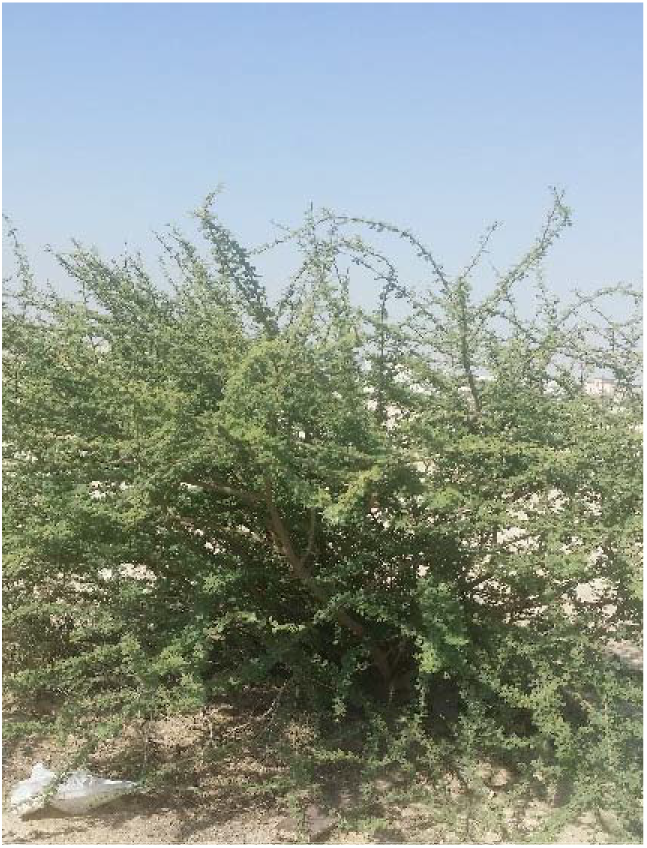
*Acacia tortilis* shrub in the study area.

The distribution of *acacia* spp. and their success in occupying certain areas depend on many factors, some of them are closely related to the physical and climate conditions particularly the wind speed and axial loads acting on the plant in compression and tension.

According to the Qatar Meteorology Department (QMD) and Qatari Civil Aviation Authority (https://qweather.gov.qa/ClimateInfo.aspx), in the end of October 2018, Qatar has suffered from heavy thunderstorms with strong wind speed that has reached 85 km/hr with maximum rainfall exceeding 84mm. Worth noting that the average rainfall in October in Doha is about 1.1mm with wind speed less than 7 km/hr. The wind and moving debris had led to the destruction of *Acacia* shrubs planted in Doha city that were originally grown under optimum conditions in Qatar Foundation Nursery. The thunderstorms accompanied by the strong winds and heavy rains have caused plants failure due to uprooting and snapping that led to breaking and felling of trees. Most nursery-grown *Acacia* plants used in gardening were damaged or broken in their weakest. On the other hand, the *Acacia* plants naturally grown in wild could persist and withstand the extreme physical conditions they were exposed to. This has stimulated us to investigate the biomechanical, morphological, and anatomical adaptations that enable wild Acacia to cope with the harsh physical conditions.

The success of a shrub in arid and semi-arid conditions depends on its adaptive capacity to the harsh physical conditions that protrude from its biomechanical properties. Young’s modulus of Elasticity, flexural stiffness, and area moment of inertia are three important parameters that are used to describe the mechanical behavior of a plant. Young’s modulus of elasticity is used to characterize the overall bending behavior of a stem regardless of its size and shape while flexural stiffness characterizes the flexibility of the stem section. The area moment of inertia rather than the diameter or the length is used to describe the geometry and the size of the plant part to which the force is applied (Hossein and Jacob, 2015). The angle of deflection is used to describe the whole stem flexibility.

The aims of the current study were to compare the biomechanical properties of *Acacia tortilis* naturally grown in wild to those grown in nursery. The current study was divided into 3 parts: 1- finding the Young’s Modulus and the Flexural Modulus of *Acacia* stem parts using materials testing machines in compression, tension, and bending, 2- Finding the deflection (bending) angle of *Acacia* stem in field using flex sensors and Arduino boards, and 3- using image processing techniques to mathematically describe the bending movement of the tree.

The biomechanical properties were studied by applying bending, compression, and tension tests using mechanical machines, flex sensors for bending, and image processing technique to simulate the stem movement and describe it mathematically. To understand the behavior and adaptations of *Acacia* to different wind speeds, its biomechanical behavior was analyzed considering its habitat characteristics.

The techniques used in this study utilize advanced technology that is being used for the first time in investigating the biomechanical properties of plants. The results of the current research are important to agricultural and mechanical engineers, botanists, gardening specialist, municipality policy makers, as well as to taxonomists and evolutionists.

Knowing the biomechanical properties of *Acacia* would enable the prediction of their growth success and failure under particular weather conditions. This is extremely important when setting plans for the conservation of rare plant species.

Analyzing the biomechanical properties of *Acacia* and generally of plant fibers would facilitate their utilization in the synthesis of synthetic polymers made of natural fibers. Using plant fibers in composite materials would reduce the pollution, the cost, and make the performance better.

Knowledge about plants biomechanics is necessary in agricultural researches aiming to stop crop loss due to stem breakage or uprooting (Robertson *et al*., 2015) as well as in forest management for the protection of large tree trunks from damage by winds (Hale *et al*., 2012). Moreover, knowledge about stem deformation when processed by machines is extremely important for materials sciences and engineering processes (Leblicq *et al*., 2015). Knowledge about plants biomechanics is also important for pure theoretical evolutionary studies as biomechanics provide with additional criteria for phenotype characterization and doing comparisons among different plant phyla and species (Shah *et al*., 2017). It also may explain the failure of certain plant species and their extinction. Biomechanics can be used to draw conclusions and suggest new hypotheses or confirm old ones discussing the evolution of land plants.

Searching the open literature has revealed that the interest in plants biomechanics is substantially increasing that is evident by the increase in the number of publications addressing the impacts of mechanical forces on plants behavior, stability, and failure (Raines *et.al.*, 2013) as this issue is critical for abstract and applied sciences as well as for commercial purposes.

Many studies had investigated the biomechanical behavior of terrestrial and aquatic plants since 1970. However, relatively few researches studied the biomechanical features of shrubs in arid and semi-arid environments (Shah *et al*., 2017).

For instance, Niklas (1997) studied the biomechanics of hollow septate internodes and concluded that increasing the internodal length decreases the bending stiffness and the ability to resist torsion. Boller and Carrington (2007) have studied the biomechanics of different seaweeds living in tidal zones that are exposed to and suffer from high wave stresses; thus, creating a high selective pressure. They concluded that only seaweeds with high flexural modulus and high ultimate tensile strength could survive, reproduce, and evolve.

Miler *et al*. (2010) have investigated the biomechanical properties of four aquatic species living in river and under danger of being washed by the stream. They concluded that the forces created by the flow of water are faced by the bending and tension abilities of the plant causing the whole system of the plant the forces affecting it to be in equilibrium. This balance holds the plants in place and is used to predict their biomechanical behavior.

Asner and Goldstein (1997) have studied the correlation between wood biomechanical properties and stem failure of trees subjected to strong winds. Their results indicated that for canopy trees under windstorms the stem condition whether it would withstand the hurricane, get uprooted or snapped is determined by its mechanical properties particularly with the elastic modulus. The snapped trees had higher elastic moduli than the other groups and that stem’s morphological characteristics such as diameter and density are not related to stem’s failure.

In their study about the physical resilience of selected shrubs in semi-arid regions, Hossein and Jacob (2015) have found that *Acacia nilotica* was the best mechanically adapted among the 5 studied species with the highest value of Young’s Modulus (332.61KPa) and flexural stiffness (7.46Nm^2^). the other 4 species were arranged in order of decreasing their mechanical properties as follows: *Ziziphus mucronate*, *Grewia bicolor*, *Acacia tortilis*, and *Bosica grandiflora* with the lowest Young’s modulus (20.94 KPa) and flexural stiffness of 2.90 Nm^2^.

Onada et al. (2009) investigated the impact of environmental conditions such as rainfall on the relationship between the biomechanical properties of the stem and the wood density in 32 plant species in Australia and concluded that the mechanical properties of plants are not merely due to genetics but also dictated by the environment.

Alvarez-Clare and Kitajima (2007) investigated the physical traits that enhance the defensive capability and survival of neotropical woody plants, and they concluded that stems with higher strength resist buckling better and suffer less from breakage due to winds and herbivores.

Gibson (2012) investigated the biomechanical properties of three plant materials (parenchyma, dense arborescent palm stems, and wood) considering their cell wall microstructure and composition. He concluded that parenchyma has the least Young’s modulus value of (0.3 MPa) and also 0.3 MPa compressive strength while the palm stem has the highest Young’s modulus value of 30 GPa and tensile strength over than 300 MPa.

## 2 MATETRIALS AND METHODS

### 2-1 Study area, samples collection and preparation

Doha Municipal is in the central-east part of Qatar with an elevation of 10m and bordered by the Arabian Gulf on its coast. Doha is covered by desert plants and shrubs with almost no trees.

The plant materials were collected in the first week of December 2018 from a partially urbanized area in Doha city called Ar-Rayyan located at latitude: 25.333883 and Longitude: 51.411684 eastern to Qatar Foundation and from Qatar Foundation Nursery (Fig. 2). A quick survey of the area and consulting the local experts revealed that *Acacia* species are abundant and are among the most dominant species in that area covering more than 25% of the total vegetation.

**Fig. 2.**
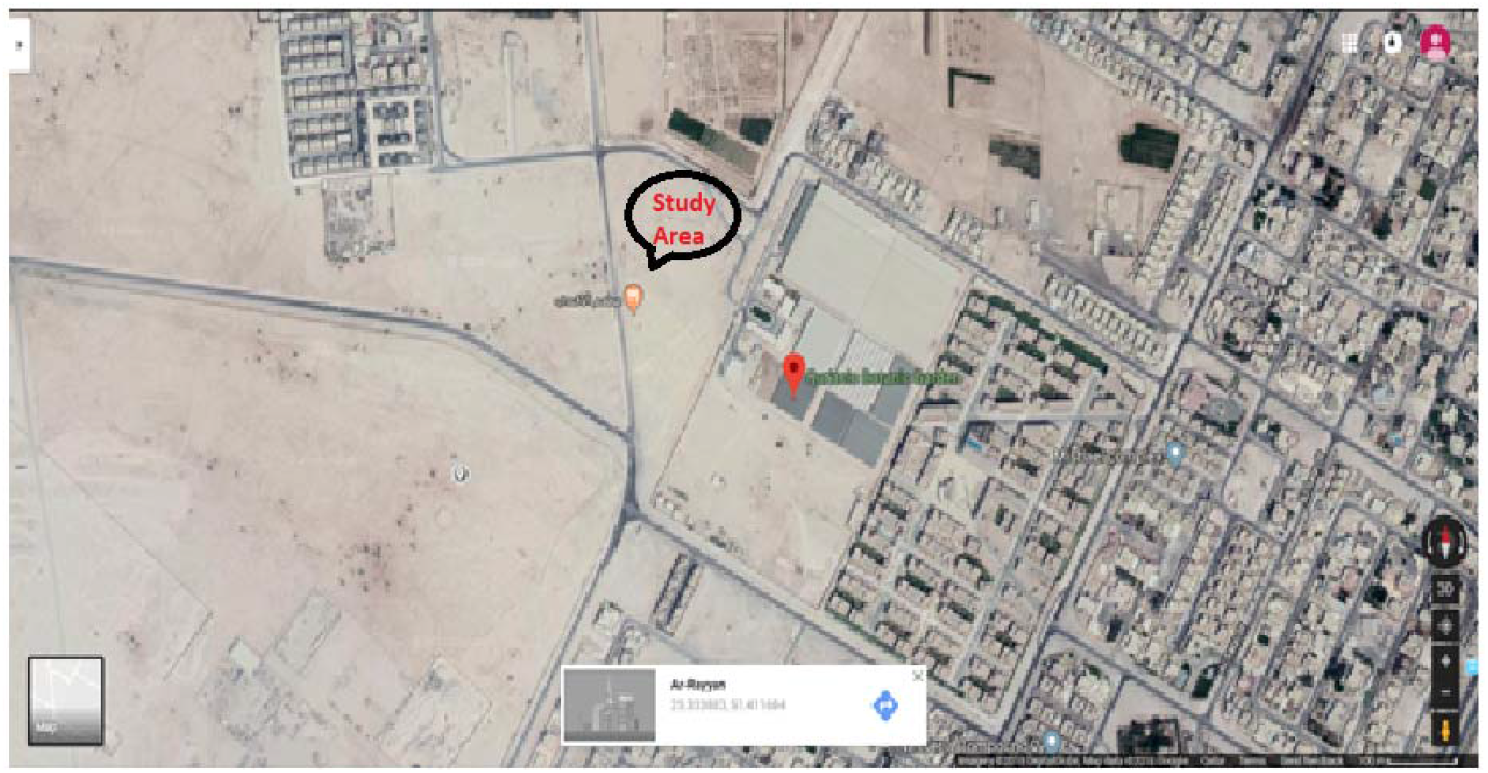
The study area showing the location of samples collection pointed at in red.

After carrying out the field study, fresh cutting specimens of *Acacia tortilis* were collected from the named location and transported to the Science, Technology, Engineering and Math (STEM) laboratories at Qatar Science and Technology Secondary School (QSTSS) for further investigations. Prior to applying the mechanical tests, the plant materials were first prepared, the thorns were removed, and initial measurements for lengths and diameters were taken using an electronic caliper. The collected samples were and stored in sealed plastic bags under cool conditions and were analyzed within one week after collection.

The length of each plant stem ranges between (50 – 85) cm. The stems were cut into three parts (top section, middle section, and bottom section), each of the sections was further cut into ∼ 10cm length, a length suitable and standardized for the bench top testing machine. The average of the length /diameter ratio was about 1:13, this ratio is enough to reduce the effect of the shear stress (Shah *et al*., 2017). For each 10 cm sample, the diameter was measured at three positions near the two ends and in the middle, or in some cases at the thickest or thinnest points, then the average value was taken.

### 2-2 Experimental setup

The collected *Acacia* samples were divided into two groups: field samples and nursery samples. For each group, the bending, tension, and compression properties were investigated using PASCO materials testing system Model ME-8230 (Fig. 3).

**Fig. 3.**
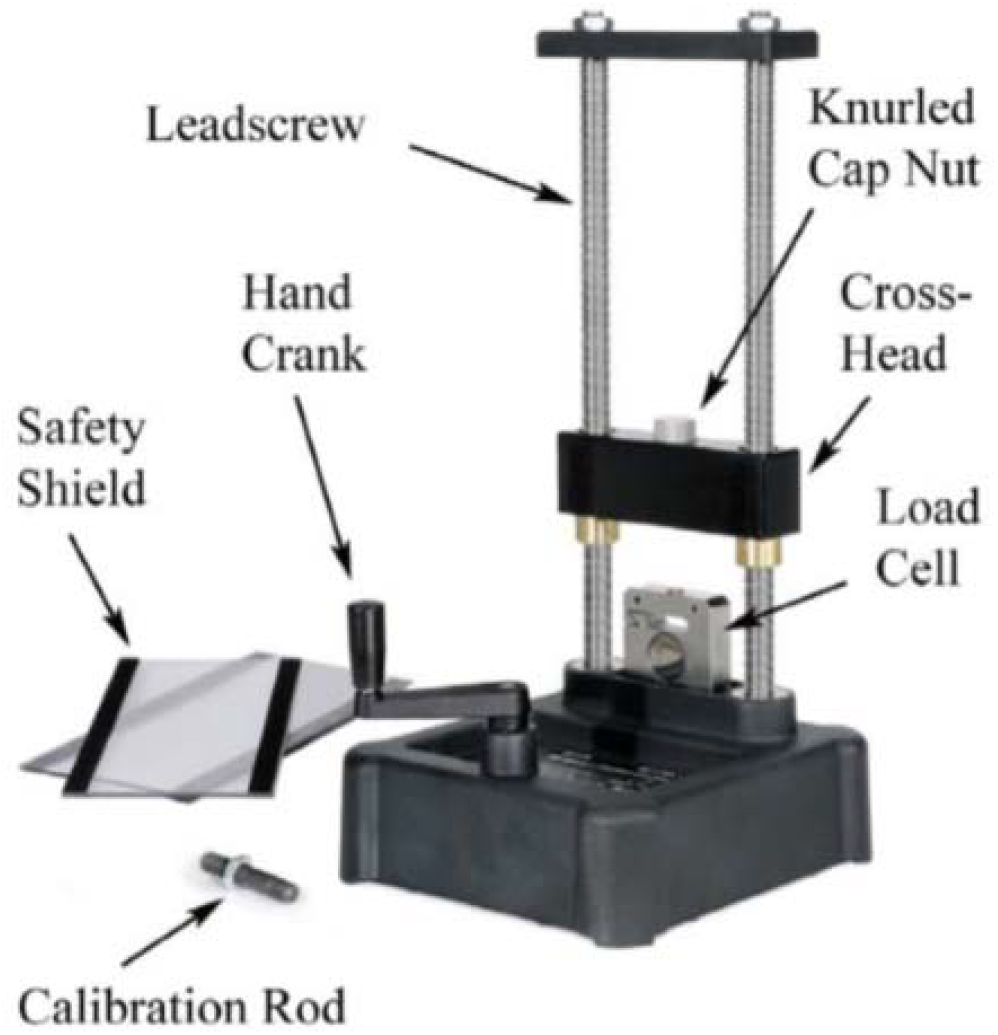
PASCO materials bench top testing machine used in the present study.

The material testing apparatus consists of two major parts: the upper load anvil and the lower base with the two support anvils. The load anvil sticks up through the cross-head and is held in place by knurled cap nut. The base (for the support anvils) fastens directly to the load cell using the two cap screws as shown in figure 3.

The materials testing machine is provided with a built-in load cell that can create and measure a force up to 7100N. The machine is also provided with a built-in optical encoder for measuring the change in the linear position (displacement) of the cross-head load bar. The load cell also contains a transducer that converts the mechanical measurements (force and displacement) into electronic signals. PASCO Interface operated by PASCO Capstone Software was used to record, save, retrieve, display, and analyze the force and displacement data from the load cell and the encoder, respectively. The materials testing machine was calibrated and used as per the instructions of the manufacturer.

Because of their anisotropic nature, taking biomechanical measurements for plant materials and interpreting them is more difficult than it is for metals. Plant tissues are heterogeneous therefore they may show different values for Young’s and Flexural Moduli when measured at different parts along the stem. Therefore, the experiment was repeated many times to determine the range of values that the moduli can take.

When carrying out the tension experiment (fig. 4), the plant material slipped from the clamps when the applied force reached a certain value so only the first portion that is the linear portion of the force vs. deflection graphs was considered valid. However, this linear portion was enough to calculate Young’s Modulus but not the tensile strength or the breaking force.

**Fig. 4.**
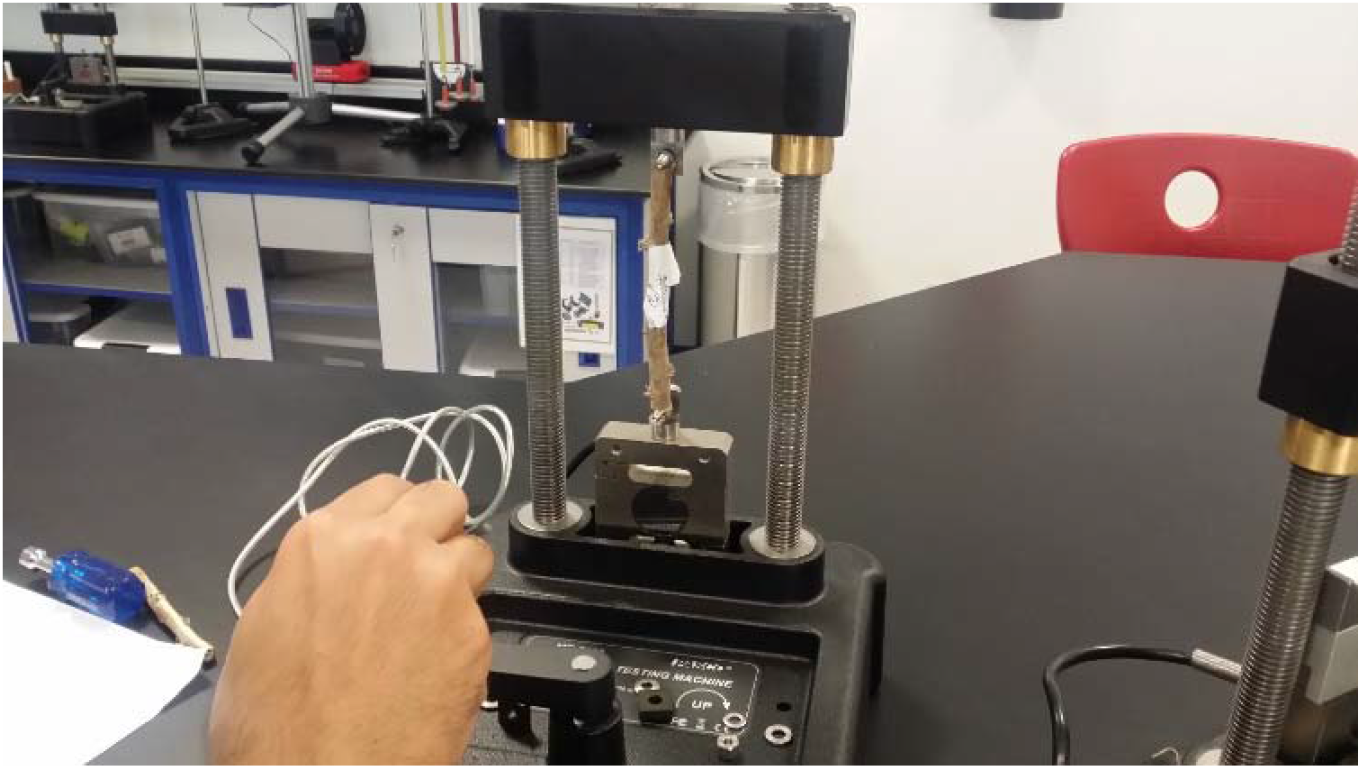
The setup for tension tests.

Force vs. deflection and stress vs. strain graphs for the plant materials within the elastic region were constructed. For each stem cut, three 10 cm samples were taken and examined. The mean value of Young’s Modulus and Flexural Modulus for the tested plant samples of the same plant cut were determined. The high accuracy of the load cells and the materials testing system used allowed us to measure the small forces during bending and compression.

### 2-3 Three Point Bending Test

Three-point bending tests were performed using a bench-top testing machine provided with two electronic sensors for measuring and recording the load force and the deflection in the form of change in position (fig. 5).

**Fig. 5.**
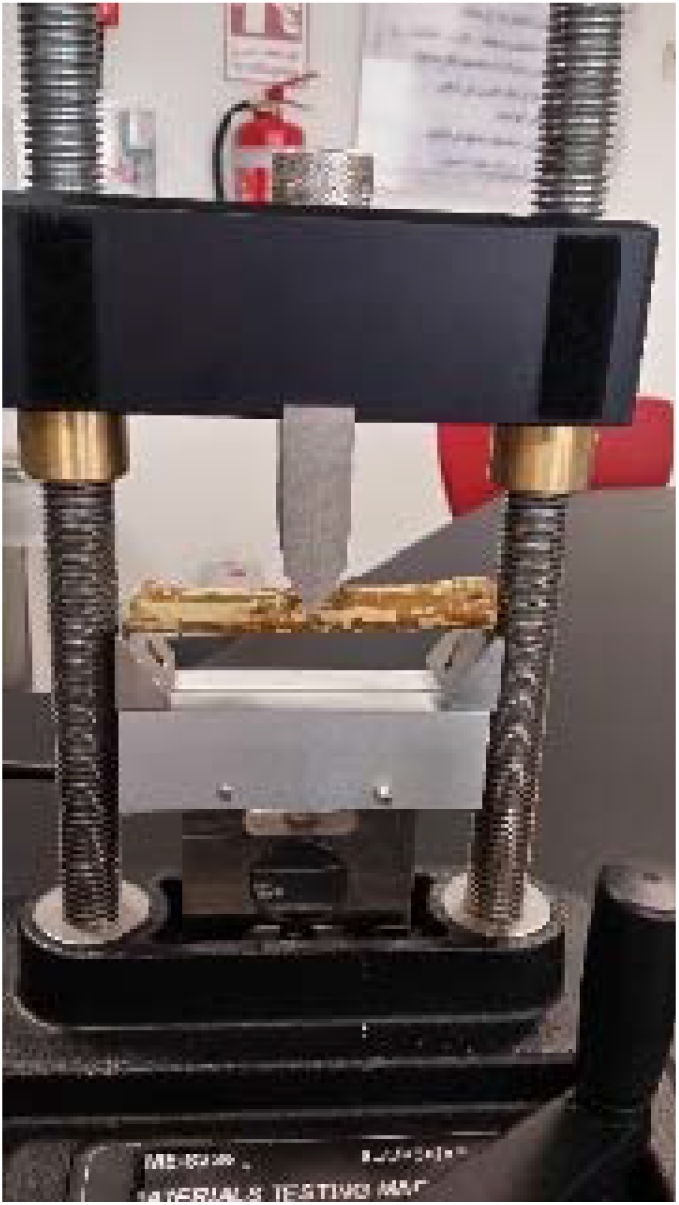
The setup for the bending test.

To apply the force on the sample through the plunger, the crank was turned slowly at a rate between 10 and 20 mm/minute. The materials testing machine was then connected to the PASCO interface for data collection and recording. The force vs. position graph was constructed.

As the plunger applied a downward load (force, F) in the middle of the plant section, the flexure (vertical deflection) was recorded. Materials tested included 20 specimens for field plant and 20 for nursery plant. For each field and nursery plant sample, the force (F, N) vs. deflection represented as a change in the linear position (ΔX, mm) was constructed and the slope was measured directly. The slope (F/ ΔX ratio) reflects the effective stiffness of the measured plant part. According to the elastic theory, the material’s stiffness depends on its length, shape and the cross-sectional area in addition to the nature of the material. The Flexural Elastic Modulus (E_b_) for the tested plant part was calculated as follows: ***F*/Δ*X* = 48*I_b_E/L*^3^**, where “**I**” is the area moment of inertia for the sample.

Solving for E_b_, yields 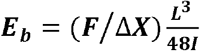.

The area moment of inertia depends on the shape of the cross section of the sample. For the round solid stem of diameter d and radius r, ***I**_stem_* = ¼ π r^4^ = 1/64 π d^4^.

The value (EI) is an indicator for the plant stiffness and important for stem length and corresponding plant height (Gere and Timoshenko, 1999).

### 2-4 Axial Loading Tests

#### 2-4-1 Tensile Tests

The plant sample (fig. 6) was firmly fixed in the materials testing machine from its two ends using two adapters that are connected to the load cell from one side and to the cross-head component from the other side. The crank of the materials testing machine was turned slowly to raise the cross-head to which the sample is attached at a rate between 10 and 20 mm/minute and pulling the stem section apart while both the load and the extension are being recorded until the stem breaks or slips.

**Fig. 6.**
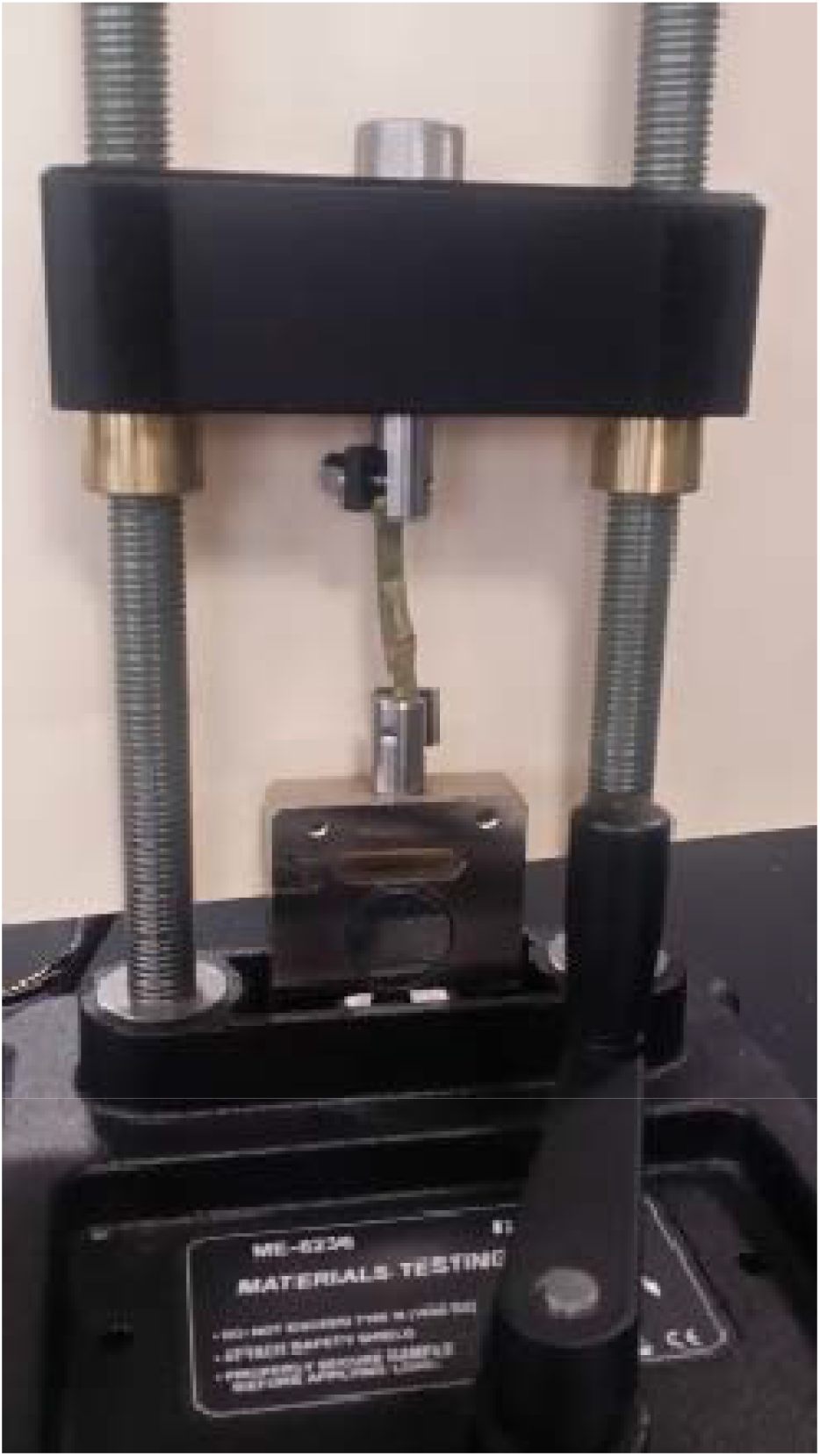
Setup for tensile tests.

The PASCO Sparkvue software was used to record the force vs displacement graph from which the breaking force was determined and the stress vs. strain graph constructed. Several different materials mechanical properties including Young’s modulus, tensile strength, resilience modulus, ductility and yield strength could be calculated from the data collected. However, in this study only Young’s modulus was calculated due to the slippage of the specimen.

The stress σ (N/m^2^) was used to express the strength of the stem section tested under tension or compression and was calculated using the formula σ = F/A, where F is the applied tensile force (N) and A is the sample’s cross-sectional area (m^2^). As the plant section is loaded under tension, it stretches and is supposed to become thinner; however, the change in diameter was neglected and the initial value of A was only considered.

The strain □ (m/m) was calculated using the formula: □ = ΔL / L, where L (m) is the plant sample’s initial length and ΔL (m) is the change in the length (amount of elongation) of the plant sample. When a plant sample is subjected to tension, it will first exhibit some temporary elongation within the elastic region then it will exhibit plastic deformation after which if the sample is pulled far enough it will break into two pieces. The maximum load that the sample can sustain is referred to as tensile strength. The stiffness of the material under tension or compression is described using Young’s modulus (E_t_ or E_c_) (N/m^2^) which is calculated from the slope of the linear portion of the stress vs. strain graph (E= σ/□). Young’s Modulus.

#### 2-4-2 Compression Buckling Test

Buckling tests help in explaining the failure of stems and determining the largest length a stem can attain and support (Frese and Blass, 2014). When an object is subjected to compression forces, it compresses until a point where the object suddenly buckles. At that point, the applied force is called the critical force (F_crit_). Euler equation (*Fcrit* = 4(*π*^2^ *EI*)/*L*^2^), where **E** is the Young’s Modulus and the **I** is the area moment of inertia, and L is the object length describes the relationship between the F_crit_ and the geometry of the object. The area moment of inertia (I) was calculated using the formula (*I_rod_* = 1/4*πr*^2^), where (r) is the radius. The buckling test (Fig. 7) was carried out as follows: the sample was first mounted in the machine. The crank was turned out slowly counter clockwise until the beam buckles. The data were recorded and the force (F, N) vs. deflection (ΔX, mm) graph was constructed. The slope (F/ΔX) was calculated from the linear portion of the graph. Young’s modulus was calculated from the formula (*E* = *L* * (*Slope*)/*A*).

**Fig. 7.**
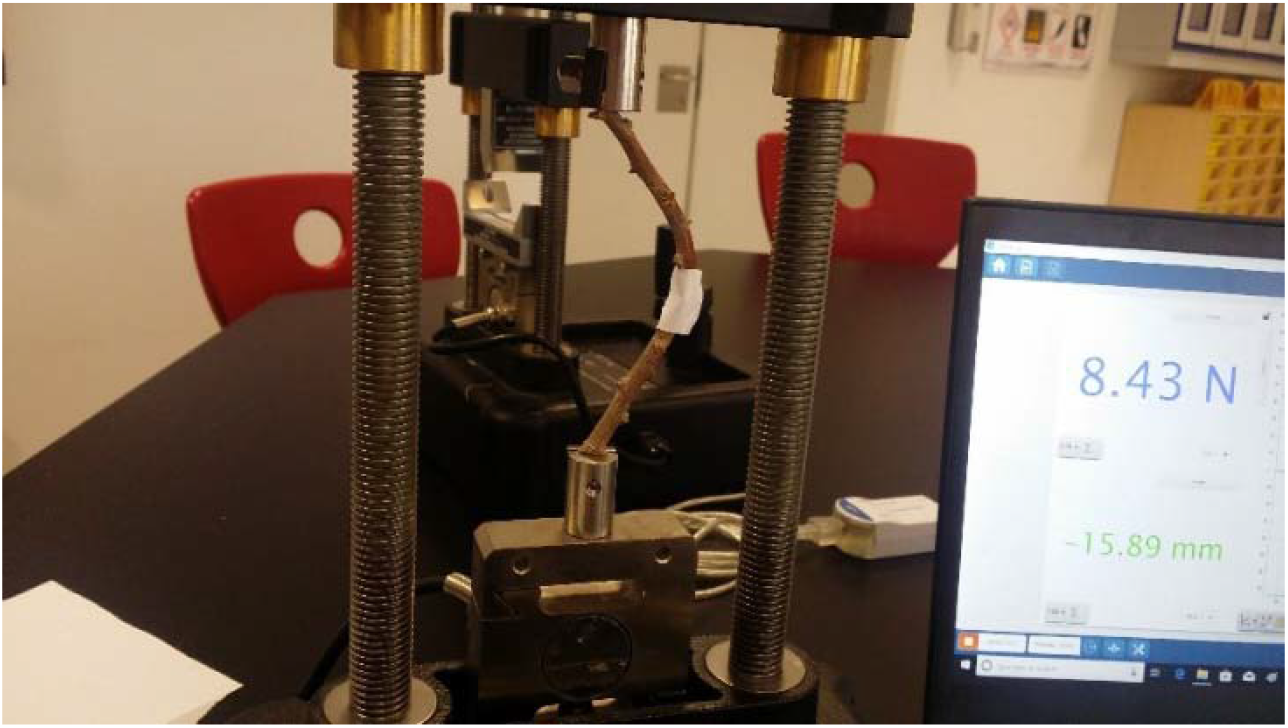
The setup for compression buckling test.

### 2-5 Deflection Angle

When a tree stem is subjected to a wind force, it bends with an angle called the deflection angle (α). The deflection angle represents the flexibility of the whole stem. It shows the bending capability of the stem section. Traditionally, the deflection angle is measured using the cantilever or two-point bending test using G-clamp to fix stem cuttings from one side and applying a known force from the free end (Shah et al., 2017, Hossein and Jacob, 2015, Caliaro et al., 2013, Moulia et al., 1994). However; in this study, the deflection angle was measured in field using flex sensors (Sparkfun.com) applied on intact living whole stem (Fig. 8).

**Fig. 8.**
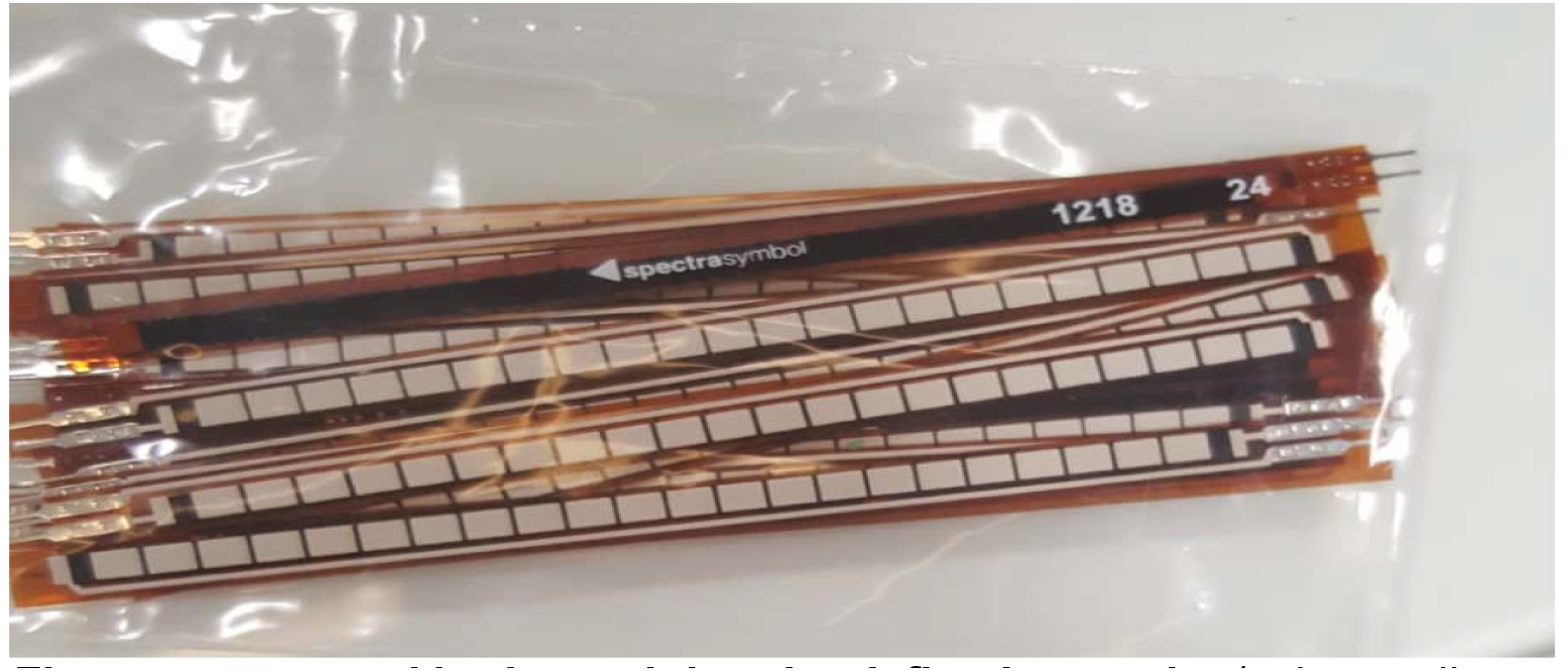
Flex sensors used in determining the deflection angle.

The Flex sensor is basically a tape shaped flexible resistor that varies with deflection. On one side, it is coated with little conductive particles. When the sensor is flat the conductive particles are close together with a total resistance of 30 kiloohms (fig. 9). The conductive particles are stretched apart when the sensor is deflected causing it to be less conductive, thus higher on resistance. The resistance was measured to be around 70 kiloohms at 90 degrees angle.

**Fig. 9.**
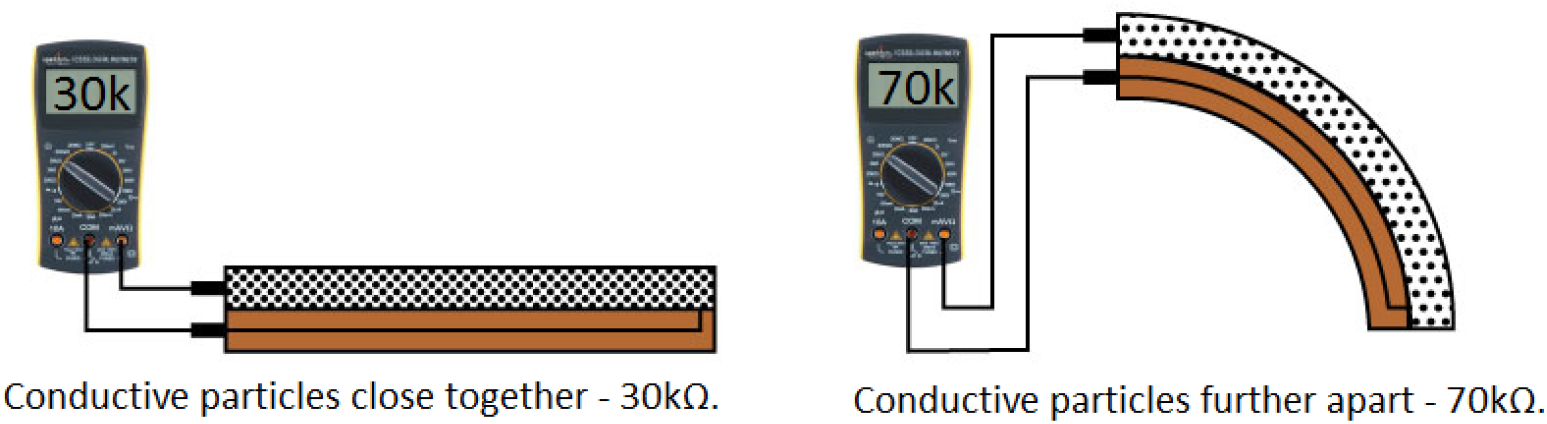
The operation of flex sensors.

Three sensors connected to an Arduino Uno board were placed along the stem (L=∼1.2m, D_nursery_=9.34mm D_field_=8.14mm) at three different positions: the base, the middle, and the end (fig. 10). Known masses ranging from 100 to 4000 grams were hanged sequentially from the smallest to the largest from the tip of the stems causing them to bend.

**Fig. 10.**
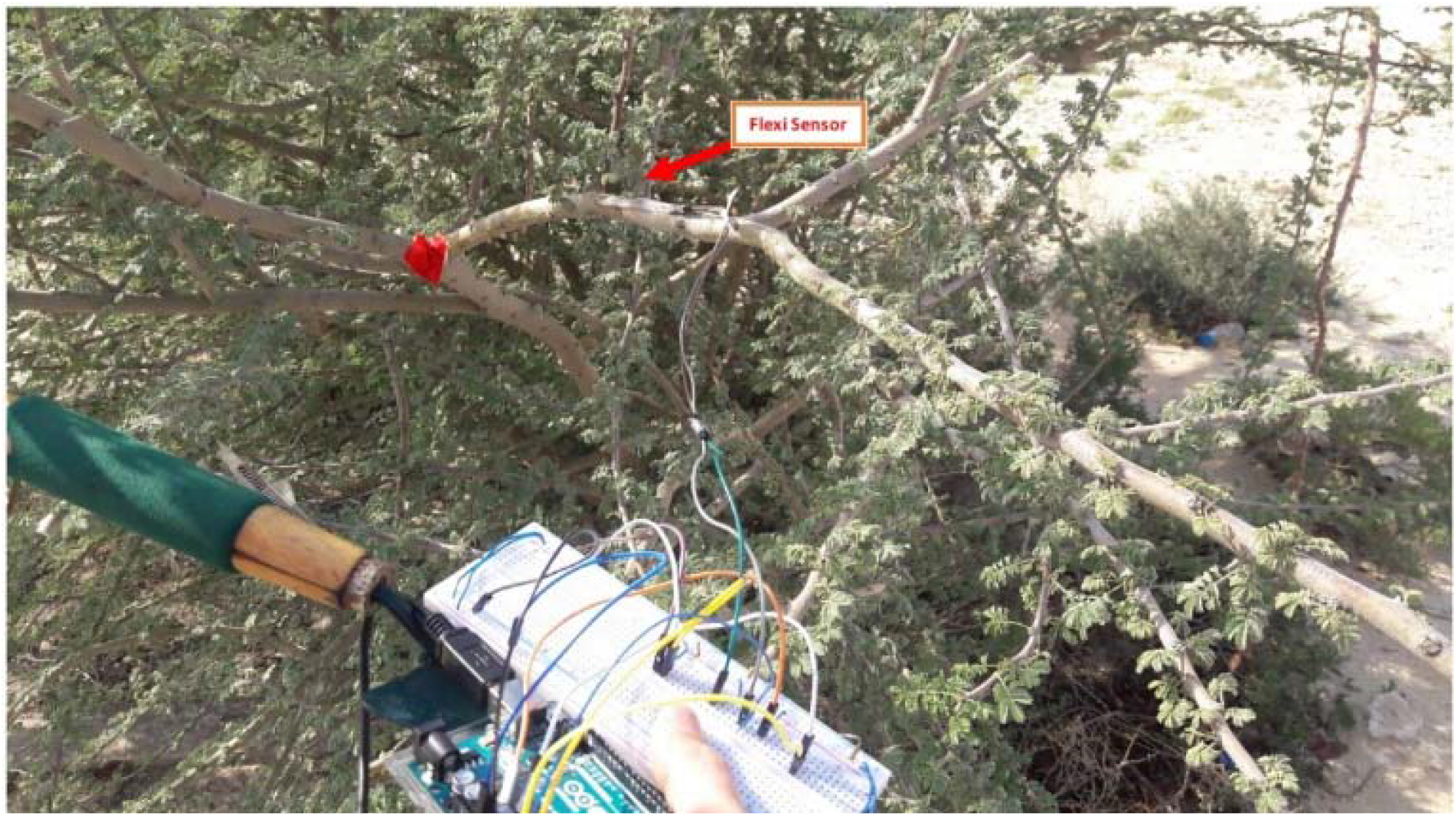
Deflection angle measurement setup.

The flex sensors detected the deflections in the tested plant part when subjected to a bending force and the Arduino board interpreted the change in resistance of the flex sensors into a change in voltage signals. The signals were sent to the laptop though serial communication and shown as angles in degrees.

### 2-6 Mathematical Modeling of the stem behavior

The Acacia branches were observed to bend as the weights hanged sequentially at their tips increase in magnitude. To find out the mathematical equation and the graph representing the branch movement when it bends due to winds or other loads, the branches were first subjected to sequential weights and the height in meters of the branch tip above the ground was measured. Estimating the mass that would cause the branch tip to reach or to touch the ground was not always a possible task as the branch length in many cases were not enough to reach the ground and in other cases it was broken before it could touch the ground. Moreover, generating the equation and the graph representing this kind of movements require enormous amount of data to ensure the accuracy and guarantee that the equation would give real solutions. However, considering the nature of the problem, we assumed that the equation would follow a polynomial function of n degree:

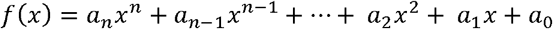

Where X represents the weights hanged from the tip and F(x) represents the height of the branch’s tip from the ground.

The equation might also be represented by a power function following the general formula:

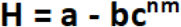

where H is the plant height at a given weight and a is the maximum plant height; the coefficient (b) and the base of the power (c) might take any real positive number; m represents the mass hanged at the tip; the coefficient (n) is a positive fraction (between 0 and 1).

In this study, image processing technique (IPT) was used to mathematically describe the actual bending movement of the tree. The continuous bending movement of *Acacia* resulting from wind forces was simulated by tying a rope to the tip of *Acacia* branch under study and then stretching the rope using known forces.

The movement of *Acacia* was tracked and videotaped. Python + Opencv were used to plot the branch motion versus time diagrams and therefore to describe the maximum forces the branch can tolerate. To do so, the plant part being investigated (branch) was covered with red tape and videotaped while subjected to a force causing it to bend (fig. 11).

**Fig. 11.**
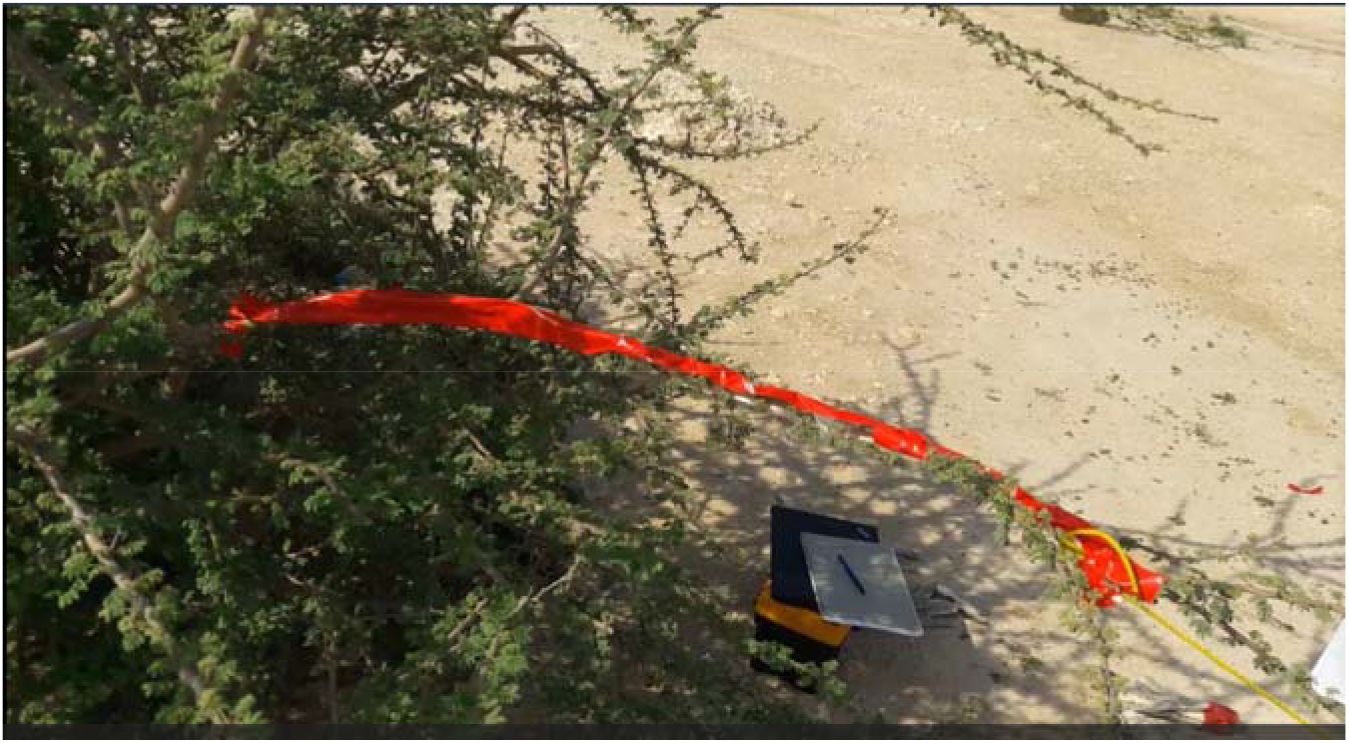
Acacia branch being studied using IPT (covered in red tape).

The video was then filtered and processed using special image processing software to highlight the bending branch. The color was converted to HSV color, and a red color filter was applied using the site (https://solarianprogrammer.com/2015/05/08/detect-red-circles-image-using-opencv/). After that binary image was produced, the red pix was changed to white pix and the rest of pixs changed to black. Then binary erosion and dilation functions were used to remove the small noise. The bending movement of the branch was then linearized and described mathematically using the site (https://scikit-learn.org/stable/auto_examples/linear_modal/plot_ols.html).

### 2-7 Statistical Analysis

T-test was performed with Excel to test if there are any statistically significant differences between the mean values of Young’s and flexural moduli between the field *Acacia* and the nursery *Acacia*.

## 3 Results

### 3-1 Bending Test

For each stem section, the length and the diameter at the midpoint and at both ends were measured then the area moment of inertia (I, m^4^) was calculated (table 1). The data show that the specimens had approximately equal length with varied diameters.

**Table 1.** Length and diameter measurements of the studied stem samples.

| Nursery Samples |  |  |  |  |  |  |  |
| --- | --- | --- | --- | --- | --- | --- | --- |
| Material Sample | D1, mm | D2, mm | D3, mm | Average D, mm | R, mm | L, m | I, m<sup>4</sup> (X10<sup>-12</sup>) |
| 1 | 15.08 | 14.82 | 14.91 | 14.94 | 7.47 | 0.095 | 2444.28 |
| 2 | 11.61 | 10.36 | 10.72 | 10.89 | 5.45 | 0.093 | 692.56 |
| 3 | 14.65 | 13.92 | 14.43 | 14.33 | 7.17 | 0.097 | 2074.66 |
| 4 | 9.77 | 9.01 | 9.23 | 9.34 | 4.67 | 0.094 | 373.37 |
| 5 | 13.79 | 13.50 | 13.70 | 13.66 | 6.83 | 0.095 | 1708 |
| Field Samples |  |  |  |  |  |  |  |
| 1 | 8.00 | 8.32 | 8.11 | 8.14 | 4.07 | 0.095 | 215.4 |
| 2 | 6.74 | 6.72 | 6.56 | 6.67 | 3.33 | 0.095 | 96.52 |
| 3 | 6.08 | 5.14 | 6.25 | 5.82 | 2.91 | 0.120 | 56.29 |
| 4 | 7.85 | 9.02 | 9.40 | 8.75 | 4.38 | 0.096 | 288.00 |
| 5 | 6.62 | 6.42 | 6.71 | 6.58 | 3.29 | 0.097 | 91.97 |

Figures 12, 13 show the force (F, N) vs. deflection (ΔX, mm) for some of the naturally grown and nursery grown *Acacia* samples subjected to bending. The slope of the linear portion (F/ΔX) represents the stiffness. The curves also show the critical force (F_crit_) after which the sample breaks.

**Fig. 12.**
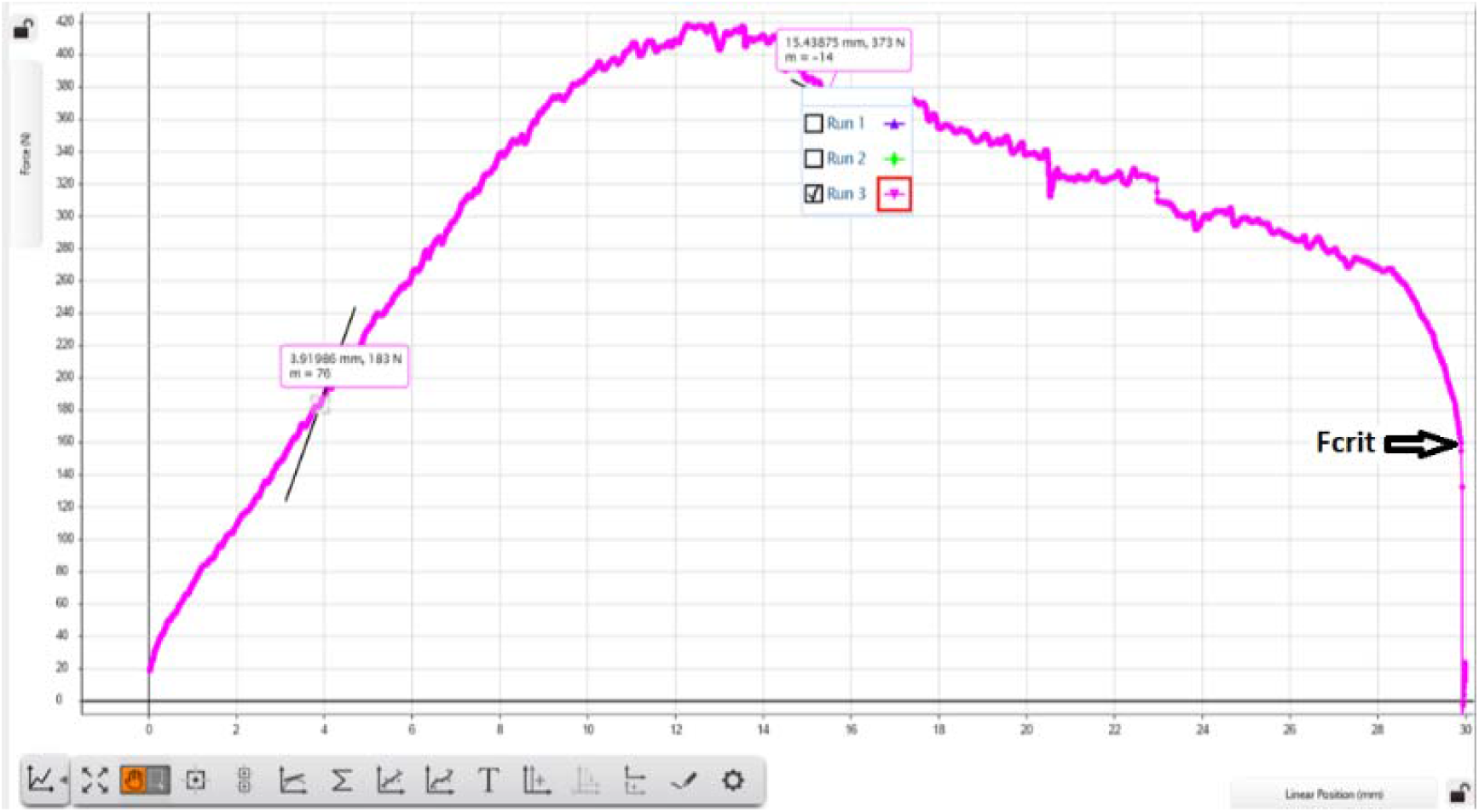
Force (N) vs. deflection (ΔX, mm) for a nursery grown *Acacia totilis*. (Slope=1.82 N/mm, F_crit_ = 160N).

**Fig. 13.**
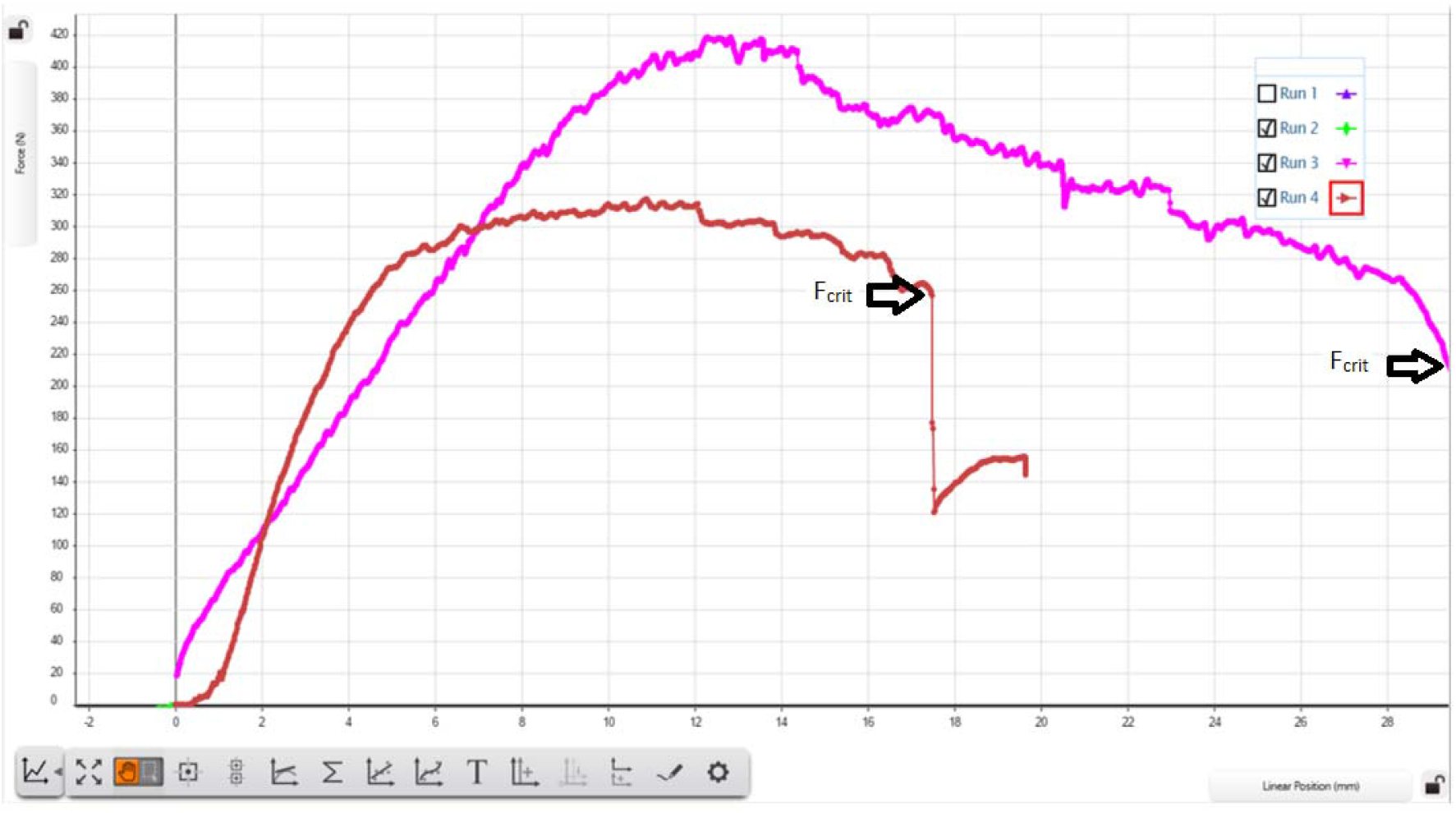
Force (N) vs. deflection (ΔX, mm) for a field grown *Acacia totilis* (brown) and a nursery grown one (violet). (Field Acacia: S=2.00 N/mm and F_crit_= 260N. Nuresry Acacia: Slope= 1.71 N/mm and F_crit_=210N).

Table 2 shows the calculations for the flexural modulus for some of the samples in the two Acacia groups.

**Table 2.** Flexural modulus values for some of the field and nursery grown Acacia specimens.

| **Three – Point Bending Test: Nursery Samples** |  |  |  |  |  |
| --- | --- | --- | --- | --- | --- |
| **Material sample** | **L, m** | **$L^3, m^3 (X 10^{-4})$** | **$I, m^4 (X10^{-12})$** | **F / $\Delta X$, (N/m)** | **$E_b$, MN/m<sup>2</sup>** |
| **1** | **0.095** | **8.57** | **2444.28** | **2000** | **14.60** |
| **2** | **0.093** | **8.04** | **692.56** | **1820** | **44.02** |
| **3** | **0.097** | **9.13** | **2074.66** | **8700** | **79.76** |
| **4** | **0.094** | **8.30** | **373.37** | **2000** | **92.62** |
| **5** | **0.095** | **8.84** | **1708** | **1710** | **18.44** |
| **Three – Point Bending Test: Field Samples** |  |  |  |  |  |
| **1** | **0.095** | **8.57** | **215.4** | **4500** | **372.99** |
| **2** | **0.095** | **8.57** | **96.52** | **2000** | **369.96** |
| **3** | **0.120** | **1.72** | **56.29** | **1820** | **1158.50** |
| **4** | **0.096** | **8.84** | **288.00** | **2000** | **127.89** |
| **5** | **0.097** | **9.13** | **91.97** | **4100** | **847.94** |

The average value of E_b_ for nursery *Acacia* was (49.88 ± 3.03, n=20) while the average E_b_ value for field *Acacia* was (575.46 ± 3.54, n=20). Since the value of the calculated t_stat_ (t=2.78) is higher than the tabulated value (t=2.13) at (P=0.05), we accept the hypothesis that states that there are statistically significant differences in the E_b_ values for field and nursery *Acacia*. The results revealed that field *Acacia* had a significantly higher value for flexural modulus than the nursery *Acacia* (table 2) indicating that the field *Acacia* is the most flexible among the two groups even though it has the smallest area moment of inertia due to the small stem diameter (table 1).

### 3-2 Axial tests: tension and compression tests

The force (F, N) vs. deflection (ΔX, m) was constructed (fig. 14). The slope (F/ ΔX) was calculated from the linear portion of the graph and was found to be 543000 N/m. The stress vs. strain graphs for the tested plant materials followed the general shape of a typical stress / strain graph starting with a linear portion then the slope gradually decreases until it flattens followed by breakage at the maximum tensile strength. Young’s modulus was calculated from the linear part of the stress vs. strain graph. At high tensile forces by the end of the test, the stem segments slipped from the clamps and no breakage occurred. Therefore, only Young’s modulus could be calculated from the slope of the linear portion of the graph before breakage and even slipping. The average Young’s modulus for field *Acacia* was found to be (617.75 ± 67.71) MPa while for nursery *Acacia* (191.25 ± 16.37) MPa. The results show a statistically significant difference between the two values (n=16, t_tab_=2.14, It_calc_I=15.37, d.f=14, P=0.05). The breaking force, breaking stress, breaking strain, and the work of fracture couldn’t be obtained for both groups (field and nursery) because at high forces the plant samples slipped from the clamps.

**Fig. 14.**
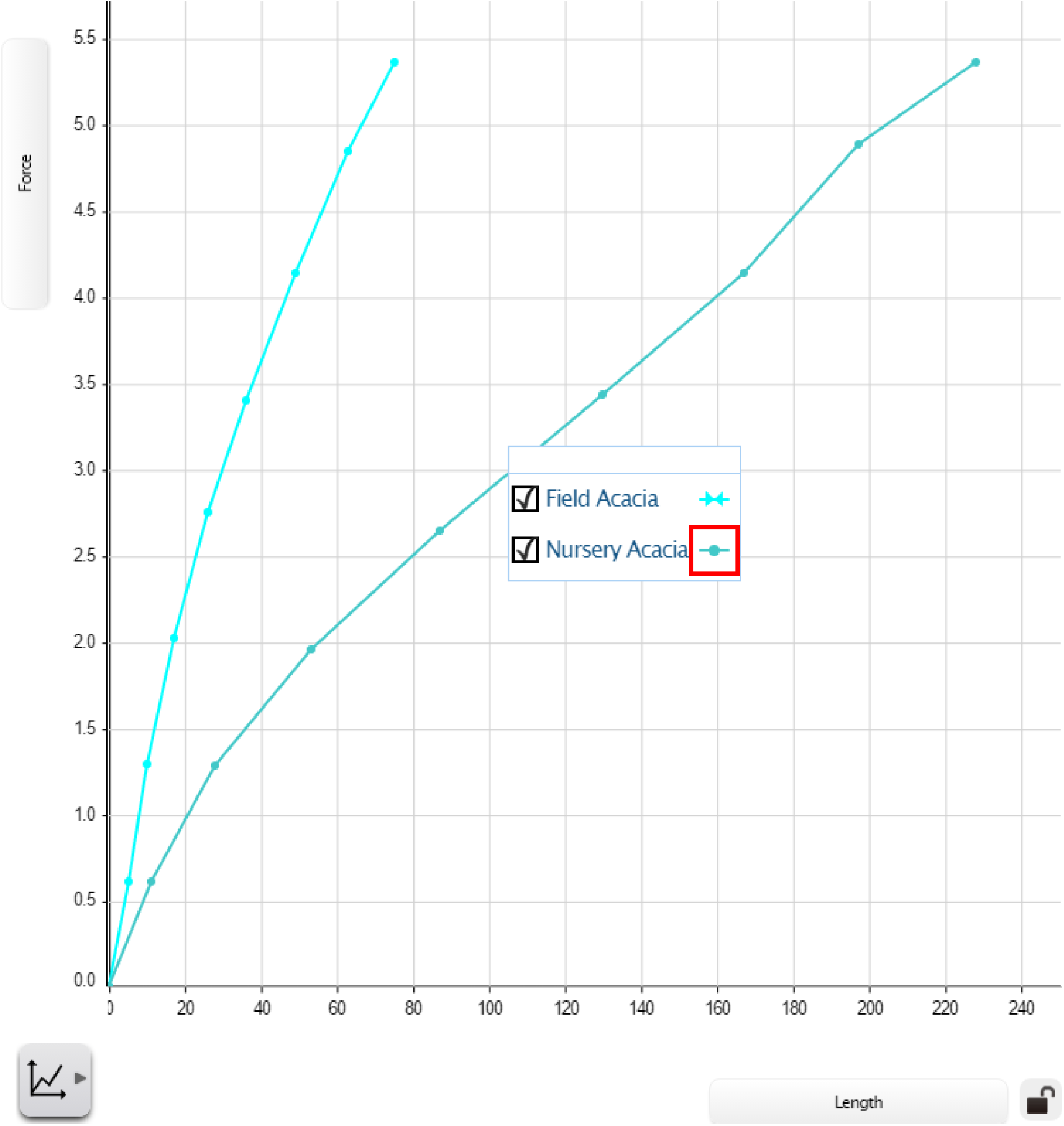
Force (F, N) vs. deflection (ΔX, m) graph for field and nursery Acacia.

All Acacia stem samples buckle when compressed. The force (F, N) vs. deflection (m) were constructed, the slope of the linear portion (F/ΔX) was calculated and the critical force (F_crit_) was determined (Fig. 15). Young’s modulus of elasticity in compression (E_c_) was calculated.

The results show that field Acacia had a significantly higher value for Young’s modulus under compression (E_c_=84.13 ± 7.77) Kpa than the nursery acacia (E_c_=17.75 ± 2.12) Kpa, (n=16, t_tab_=2.14, t_calc_=32.3, d.f=14, P=0.05) (table 3).

**Fig. 15.**
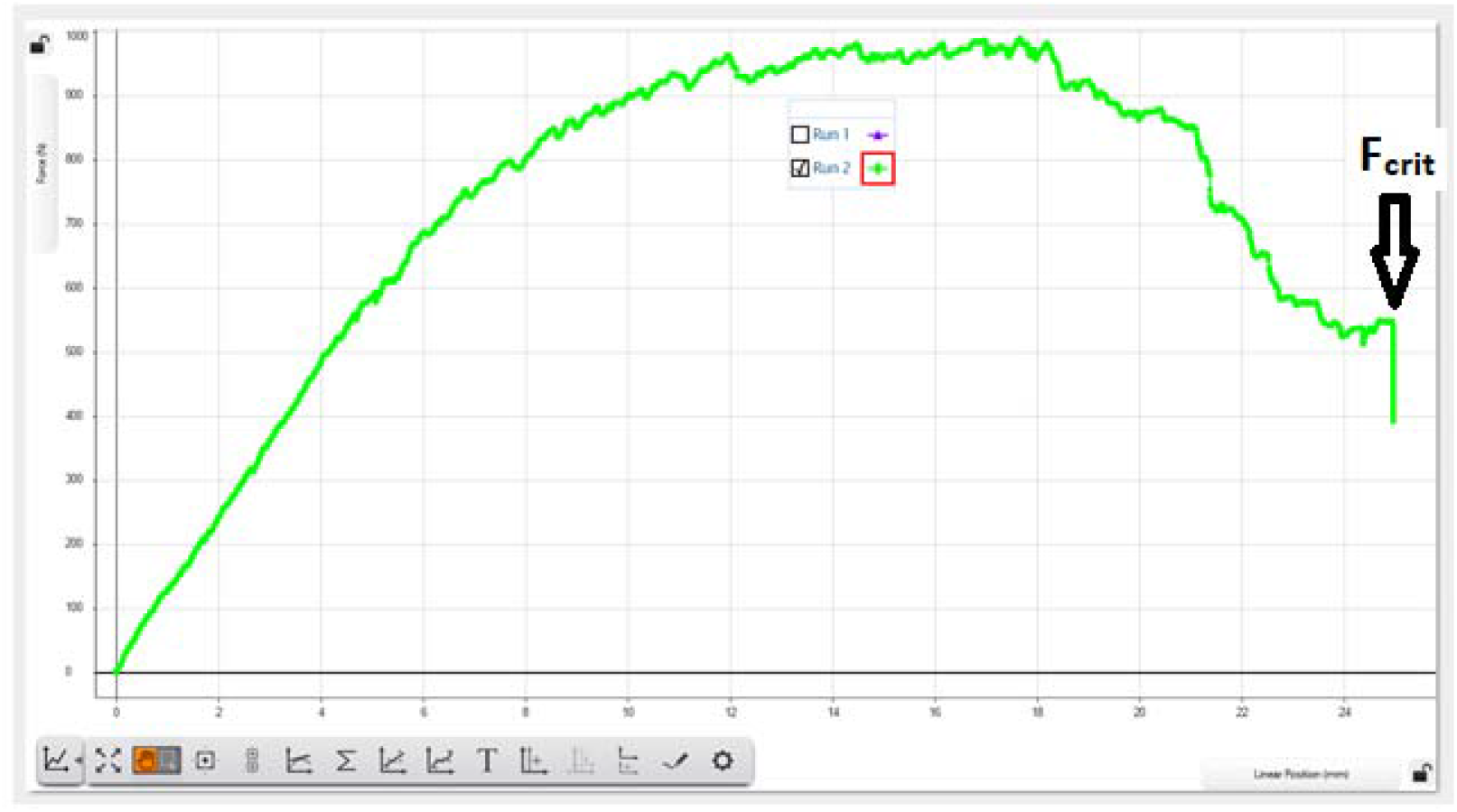
Force (N) vs. deflection (ΔX, mm) for a field grown *Acacia totilis*. (S=2 N/mm, F_crit_ = 550N).

**Table 3.** Mean ± S.D. values of the biomechanical properties of the studied field and nursery Acacia groups (n=20).

| Biomechanical Parameter | Unit | Mean |  |
| --- | --- | --- | --- |
|  |  | Nursery Acacia | Field Acacia |
| Mean Radius | mm | 6.32 | 3.59 |
| Mean C.S. Area | mm<sup>2</sup> | 125.41 | 40.46 |
| Young's Modulus ($E_t$) | MN / m<sup>2</sup> | 191.25 ± 16.37 | 617.75 ± 67.71 |
| Young's Modulus ($E_c$) | KPa | 17.75 ± 2.12 | 84.13 ± 7.77 |
| Breaking force ($F_{crit}$) | N | 210 | 550 |
| Flexural Modulus ($E_b$) | MN / m<sup>2</sup> | 49.88 ± 3.03 | 575.46 ± 3.54 |
| Area Moment of Inertia | m<sup>4</sup> | 1458.58 | 149.64 |
| Flexural rigidity ($E_b I$) | MNm<sup>2</sup> | 72753.97 | 86111.24 |

### 3-3 Deflection Angle

The results (table, 4) show that the maximum deflection (bending) angle of *Acacia* grown in field is approximately 50∘ (fig. 16) and is greater than that of *Acacia* grown in nursery. Therefore, it was concluded that the field Acacia is more elastic than the nursery Acacia as the higher the angle, the higher the elasticity.

**Table 4.** Whole stem flexibility values (force / deflection angle) for field and nursery Acacia.

| **Mass, g** | **F, N** | **Deflection**<br>**Angle**<br>**(Field)** | **Ratio**<br>**(Force/Angle)** | **Deflection**<br>**Angle**<br>**(nursery)** | **Ratio**<br>**(Force/Angle)** |
| --- | --- | --- | --- | --- | --- |
| **100** | **0.98** | **7** | **0.14** | **8** | **0.12** |
| **500** | **4.9** | **15** | **0.37** | **18** | **0.27** |
| **1500** | **14.7** | **22** | **0.67** | **28** | **0.53** |
| **2000** | **19.6** | **28** | **0.70** | **32** | **0.61** |
| **2500** | **24.5** | **37** | **0.66** | **38** | **0.64** |
| **3000** | **29.4** | **43** | **0.68** | **breakage** |  |
| **4000** | **39.2** | **50** | **0.78** |  |  |

**Fig 16.**
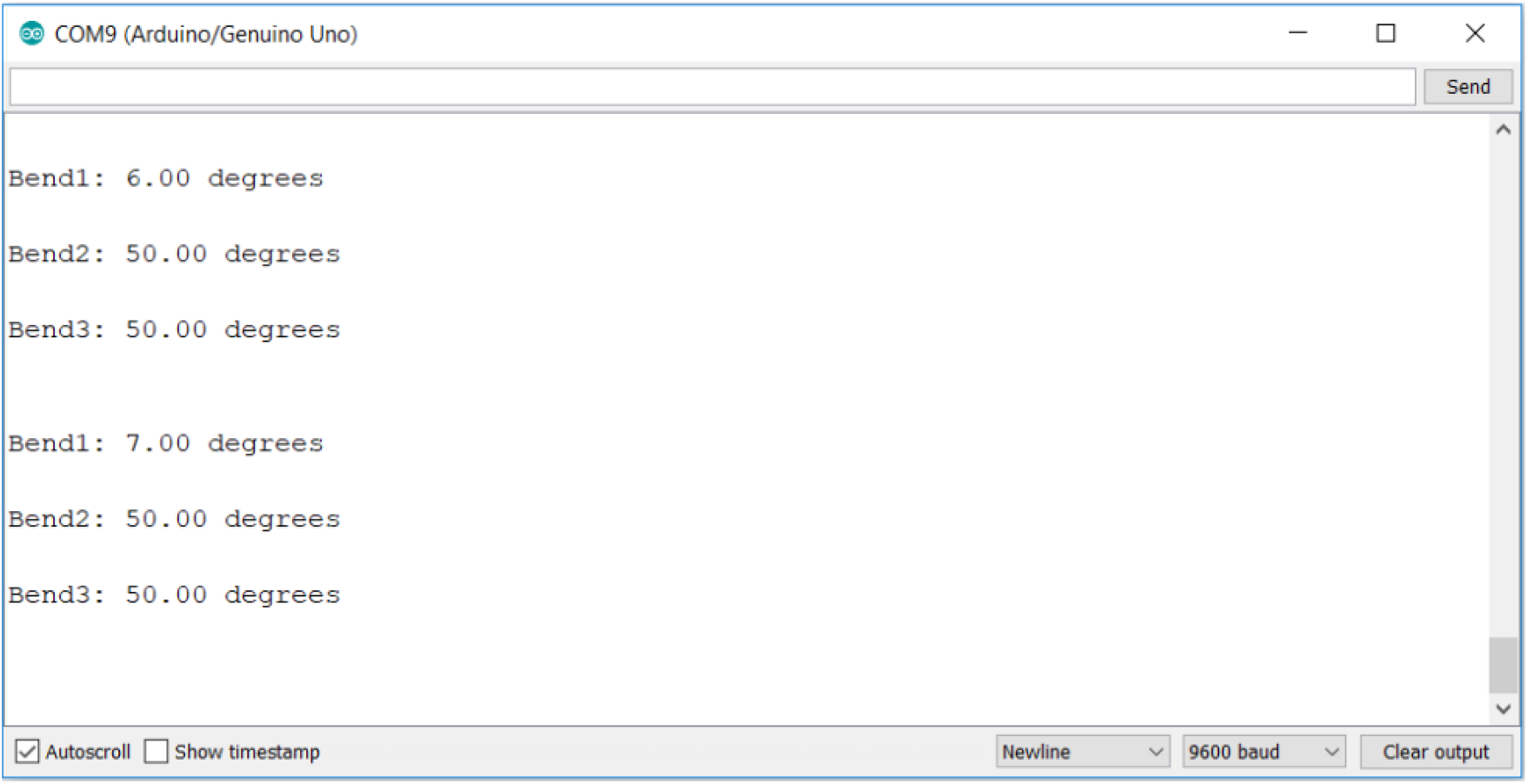
Snapshot of the computer screen showing the output of the Arduino IDE (www.arduino.cc) expressed in deflection angles.

### 3-4 Mathematical modeling of the stem’s movement during bending

(0.1-4) masses kg were hanged from the tip of the branch and its height from the ground was measured (table 5).

**Table 5.** Change in the branch tip height from the ground in response to the hanged masses.

| **Mass, Kg** | **Force, N** | **Branch tip height**<br>**above the ground, m** |
| --- | --- | --- |
| **0** | **0** | **1.2** |
| **0.1** | **0.98** | **0.91** |
| **0.5** | **4.9** | **0.73** |
| **1.5** | **14.7** | **0.66** |
| **2** | **19.6** | **0.52** |
| **2.5** | **24.5** | **0.4** |
| **3** | **29.4** | **0.32** |
| **4** | **39.2** | **0.18** |

Applying the regression analysis using the software (http://www.xuru.org/rt/), it was revealed that the data follow the binomial equation:

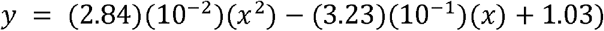

Residual Sum of Squares: rss = (6.11)(10^−2^),

Coefficient of Determination: R^2^ = (9.21)(10^−1^),

Where y is the branch tip height above the ground. However, there are no real solutions for this equation at (y = 0) making the above equation not useful in finding the force that can cause the branch tip to touch the ground. Moreover, the limited number of data makes the probability of errors high. Therefore, we decided to use image processing techniques to get a more accurate equation based on the observation that when the *Acacia* branch is subjected to a bending force, it moves in a parabolic fashion. To mathematically represent this movement in space with time, the stem was divided into 745 successive points each has its own spatial coordinates based on a given reference system (fig. 17). As time passes with the movement of the stem, the coordinates of the successive points change yielding an overall parabolic diagram. Image processing technique (IPT) was used to determine the initial and the final position for each of the points. Then the change in position with time taken frame by frame was converted into a change in position expressed through mathematical parameters (table 6) using the site (https://scikit-learn.org/stable/auto_examples/linear_modal/plot_ols.html) (Pedregosa *et al*., 2011).

**Fig. 17.**
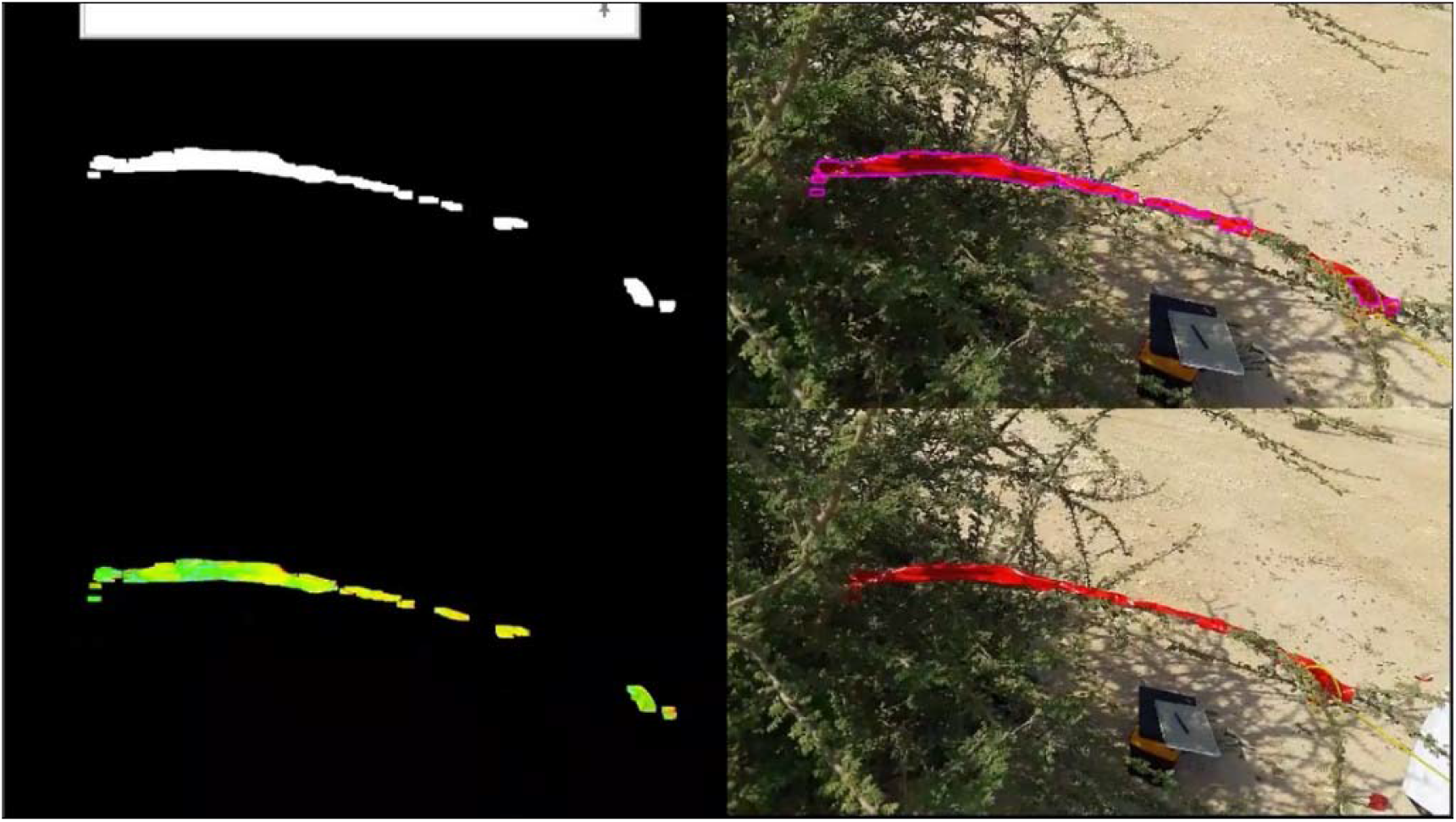
A snapshot of the video illustrating the results of IPT. The lower right panel shows the branch under study covered in red tip while subjected to a bending force. The right upper panel shows the branch highlighted. The left panels show the branch after the color was converted to HSV and a red filter was applied to produce a binary image. The red pix was changed to white and the rest of pixs were changed to black. To watch the video, follow the link (https://youtu.be/TSjYAzsahTk).

**Table 6.** The slope, Y-intercept, r-value, p value and st. dev. for the first 10 points.

| Point No. | slope | intercept | r-value | p-value | stderr |
| --- | --- | --- | --- | --- | --- |
| 1 | 4.453618 | -1635.6 | 0.495481 | 0 | 0.045093 |
| 2 | 4.589312 | -1712.43 | 0.523913 | 0 | 0.043236 |
| 3 | 4.76606 | -1798.82 | 0.554214 | 0 | 0.040997 |
| 4 | 4.645092 | -1744.36 | 0.551811 | 0 | 0.04082 |
| 5 | 4.750322 | -1790.95 | 0.562888 | 0 | 0.040882 |
| 6 | 5.131369 | -1985.66 | 0.618514 | 0 | 0.038065 |
| 7 | 4.420355 | -1619.58 | 0.521333 | 0 | 0.042286 |
| 8 | 4.396112 | -1604.37 | 0.530569 | 0 | 0.040691 |
| 9 | 4.031862 | -1416.58 | 0.488447 | 0 | 0.041636 |
| 10 | 4.39763 | -1614.49 | 0.560166 | 0 | 0.037358 |

In table 6, the slope, the Y-intercept, the r value, the P value, and the standard deviation describe the linear equations of the lines connecting the initial and the final positions of the 745 points. Knowing the equations of the lines joining the initial and the final positions due to a certain force can help determining the polynomial or the parabolic equation for the stem branch in its initial rest state and its final state as a function of force; thus, enable making predictions about the movement due to a variable force causing bending.

Using the abovementioned software, it was shown that the data follow the equation:

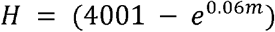

Where H is the height of the plant tip above the ground and m is the mass of the load in Kg (fig. 18).

**Fig. 18.**
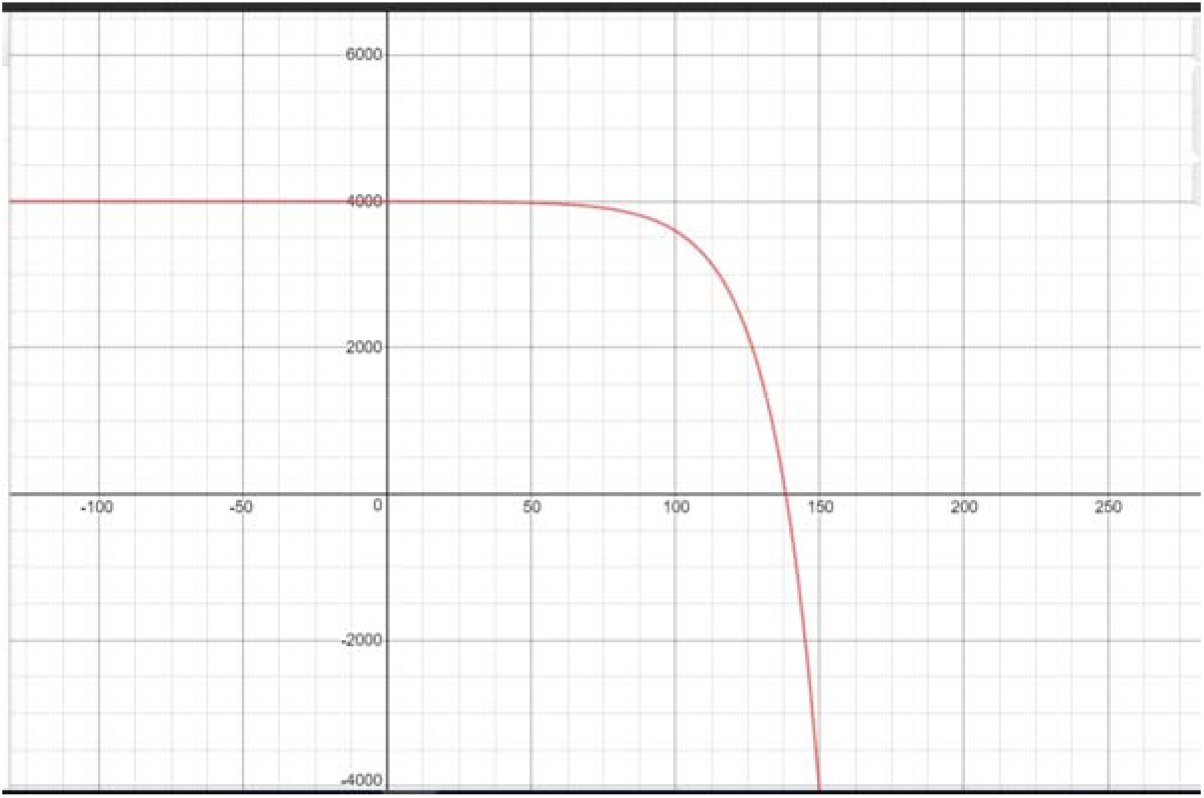
The graph representing the movement of the stem when subjected to different bending forces.

## 4 Discussion

Knowing the biomechanical properties of *Acacia* enables the prediction of their growth success and failure under particular weather conditions. Searching the open literature revealed that there no studies related to plants biomechanics in Qatar. This topic, plants biomechanics has never been addressed before in this region. In the current research, the mechano-morphological properties of two acacia groups naturally grown in field and nursery grown were investigated and analyzed for the first time. The plant – habitat interaction was investigated in this study to determine the way in which *Acacia tortilis* responds mechanically in controlled (nursery) and dry (field) environments. All measurements and calculations based on Young’s modulus of elasticity and flexural modulus were taken on stems at a nearly equal diameters in order to reduce any effects of size on the values of the measured biomechanical properties.

The study was divided into 3 parts:

1- Using mechanical instruments to find out Young’s and Flexural moduli:

Using three-point bending test and axial tests, it was found that naturally grown *Acacia* in fields have better mechanical properties than nursery grown ones. This explains the local wide spread of *Acacia* and their ability to resist harsh environmental conditions such as rain beating and strong winds where nursery grown *Acacia* failed to do. It seems that growing *Acacia* under special conditions in nursery in abundance of nutrients and water, would increase the plant’s biomass, resulting in increase in the number or size of the parenchyma storing cells. Those cells have primary thin cell walls and therefore less amount of the mechanically outstanding polymer “cellulose”. On the other hand, naturally grown *Acacia* in poor soil with limited amount of water results in the reduction in the size and / or the number of the mechanically weak parenchyma cells. This gives more space for sclerenchyma and collenchyma cells to grow. These cells have thicker cell walls with more amount of cellulose, hemicellulose, lignin, and pectin. Moreover, field *Acacia* has developed stronger hard wood due to well-developed xylem vessels. However, to verify this result, cross sections in field and nursery *Acacia* must be prepared and compared.

*Acacia* grown in nursery with high soil nutrients and good irrigation tend to have less dense stem compared to *Acacia* grown in field in dry and low nutrient soil. The density is correlated with the biomechanical properties of the stem. We recommend measuring the density as the shrub’s density gives an indication for the cellulose and hemicellulose fibers content in the cell. The bark in *Acacia* stems has low strength and removing it would not affect the overall biomechanical behavior of *Acacia* (Fig. 19).

**Fig. 19.**
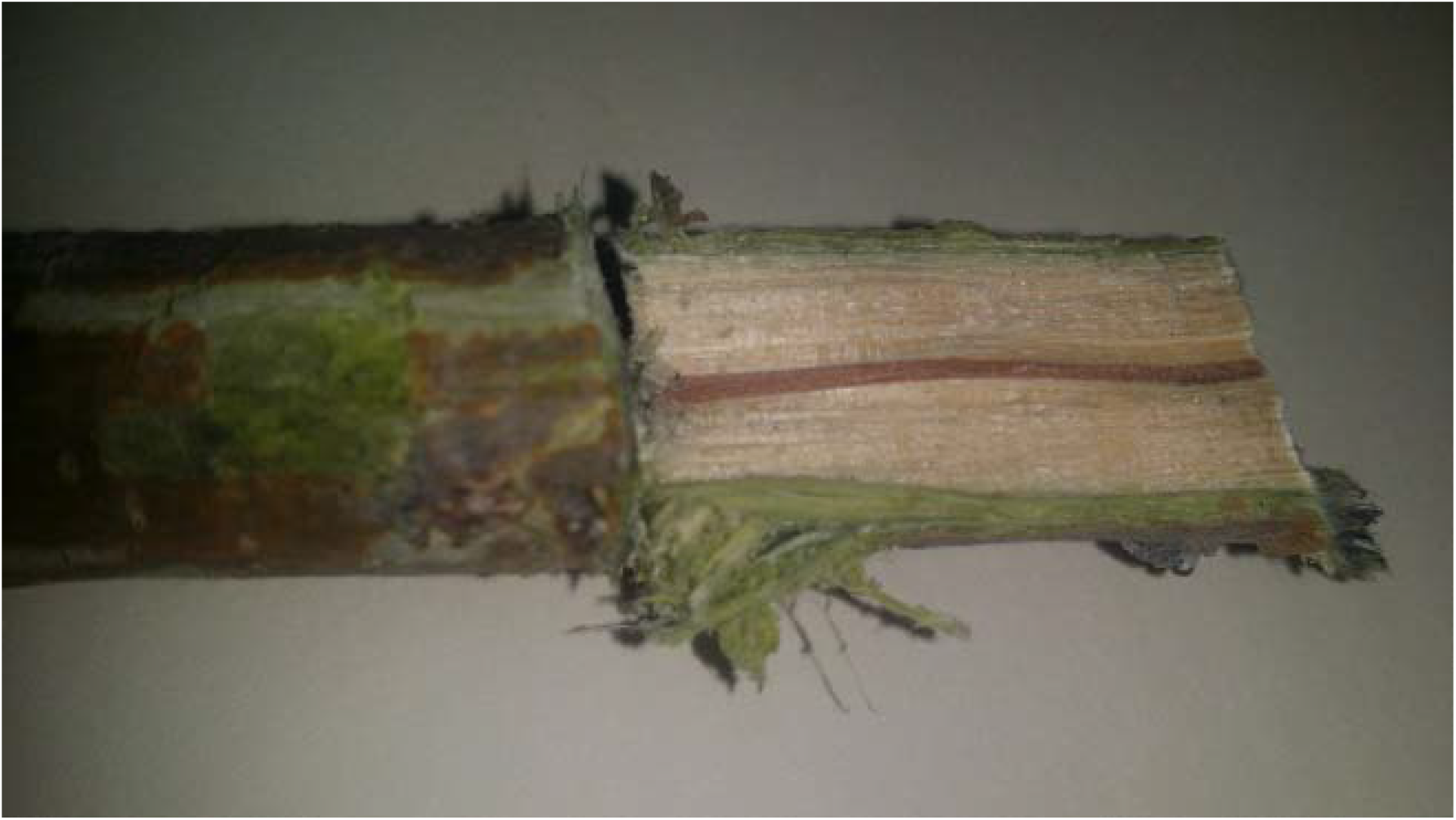
Longitudinal section of field Acacia showing the bark and the hard wood.

Among the biomechanical features that make a shrub competent in its environment are Young’s Modulus of Elasticity that describes the overall bending properties of a stem despite of the size and shape. Our values for flexural modulus generally match the results of other studies investigating the biomechanical properties of *Acacia* or a similar shrub including the study of Nakai T. (1985), Rokeya *et al*. (2010), Hossein and Jacob (2015), and the wood database (https://www.wood-database.com/koa/). However, our value for Young’s Modulus of Elasticity is less than the reported values in the above studies. This can be partially explained by the conditions of the test e.g. how moist or fresh was the stem at the time of testing or to the differences in the accuracy of the different machines and instruments used. Shah *et al*. (2017) in their review about the theory and experiments used in studying plants strength thoroughly discussed the pros and cons of each of the mechanical tests and the proper conditions for each test. It was also noted that the flexural modulus and the Young’s modulus have different values for all the studied samples. While axial forces result in either compressive or tensile stresses, bend testing produces both tensile and compressive stresses.

2- Using electronic sensors to find out the angle of deflection:

Stem angle of deflection is an indicator for the bending capability of the whole stem and for stem’s sections. Traditionally, deflection angles are measured in the lab using cantilever bending test by fixing the stem cutting to the bench using G-clamp and tying loads to the free end; however, in this study the deflection angles were measured in field using flex sensors.

The results showed that naturally grown field *Acacia* have larger deflection angles than the nursery grown ones. Having large deflection angles implies that the field *Acacia tortilis* can withstand strong winds and loads by bending without being broken. However, the nursery *Acacia* with their low deflection angles are more susceptible to breakage under strong winds.

The whole stem flexibility was calculated as a ratio between the deflection angle and the weight applied. The results show that field *Acacia* has higher value than nursery *Acacia* and this must to a certain extent reflect the survival value of both shrubs.

3- Using image processing techniques to mathematically describe the motion of the Acacia stem when bending.

It was shown how image processing technique along with python and opencv can be used to describe the movement of the bending stem as a displacement time function. The stem was divided into successive points, and when it bends the position in space of these points change. The equations of the lines connecting the initial and final positions of these points were described using the slope, the Y-intercept, the r- and the P values.

This work can be extended to make predictions about the behavior of the *Acacia* stem under any variable forces causing bending. Predicting the bending behavior of *Acacia*, can help in planting it in a manner that makes it less suffer from beating rains and strong winds.

*Acacia tortilis* grown in arid wild conditions develop many thin stems, each bear thinner branches with reduced size but densely arranged leaves. The height of *Acacia* reaches 160cm with about 7cm diameter. The thin *Acacia* branches intermingle with each other forming a web like structure. This plant configuration enables it to withstand strong wind speeds as it reduces the pressure created by axial and orthogonal forces (stresses) acting on it by increasing the overall surface area. The plant network configuration makes the thin branches connected to each other via hinge (pivot) or node like connections, so if any of the branches is dragged or pulled, other branches will be pulled too. This way of pressure transmission and distribution over a wide plant surface area reduces the stress yielded by wind, sand, beating rain, snow and confers the plant structure more elasticity under bending, tension and compression conditions. However, for this to be achieved *Acacia* must possess high level of flexibility and elasticity.

The biomechanical properties of *Acacia* shoots enable it to reconfigure and adapt to the harsh sandy winds and rainy storms. Therefore, when *Acacia* plants are planted or trimmed much attention must be paid to their biomechanical properties otherwise failure would be unavoidable. The overall surface area and the number of main stems bearing the overall plant’s weight determine the plant’s success or failure. Hence, the proper growth and distribution of *Acacia* would make them potent wind breaks and confer them economic importance.

## Acknowledgement

We would especially like to thank every member in the school administration particularly the QSTSS principal, Mr. Mohammed Al-Emadi who supported us during the entire course of the research. Big thanks to Mr. Ahmad Alghareib from Qatar Foundation for his help with the identification of the plant. We would also like to express our deepest gratitude to Mr. Yamen Farah and Mr. Muhammad Qasim from QSTSS for their help in working the math part in this study.

